# Slc9a6 mutation causes Purkinje cell loss and ataxia in the *shaker* rat

**DOI:** 10.1101/2022.03.28.486143

**Authors:** Karla Figueroa, Collin J. Anderson, Sharan Paul, Warunee Dansithong, Mandi Gandelman, Daniel R. Scoles, Stefan M. Pulst

**Affiliations:** University of Utah Department of Neurology, Salt Lake City, UT

**Keywords:** Ataxia, Cerebellum, Slc9a6, NHE6, Shaker rat, Christianson syndrome, AAV

## Abstract

**Background:** The *shaker* rat carries a naturally occurring mutation leading to progressive ataxia characterized by Purkinje cell (PC) loss. We previously reported on fine-mapping the shaker locus to the long arm of the rat X chromosome. In this work, we sought to identify the mutated gene underlying the *shaker* phenotype and confirm its identity by functional complementation.

**Methods:** We fine-mapped the candidate region and analyzed cerebellar transcriptomes to identify deleterious variants. We generated an adeno-associated virus (AAV) targeting solute carrier family 9, member A6 (*Slc9a6)* expression to PCs using a mouse L7-6 (L7) promoter, as well as a control green fluorescent protein (GFP)-expressing virus. We administered AAVs prior to the onset of PC degeneration through intracerebroventricular injection and evaluated the molecular, cellular, and motor phenotypes.

**Results:** We identified a XM_217630.9 (*Slc9a6*):c.[191_195delinsA] variant in the *Slc9a6* gene that segregated with disease. This mutation is predicted to generate a truncated sodium-hydrogen exchanger 6 (NHE6) protein, p.(Ala64Glufs*23). Administration of AAV9-PHP.eB expressing rat *Slc9a6* prior to symptom onset reduced the *shaker* motor, molecular, and cellular phenotypes.

**Interpretation:** *Slc9a6* is mutated in *shaker* and also in human Christianson syndrome, an epileptic encephalopathy. AAV-based gene therapy may be a viable therapeutic strategy for Christianson syndrome, and the *shaker* rat model may aid in therapeutic development.

## Introduction

Degenerative cerebellar ataxias comprise a large and heterogeneous group of disorders. Symptoms, age of onset, rate of progression, and prevalence are highly variable. At least in part due to this heterogeneity, symptoms are difficult to treat, and most patients never experience more than symptomatic care. Animal models specific to individual forms of ataxias may ultimately prove important in generating new therapeutic strategies. While there are a number of animal models of ataxia, particularly for the spinocerebellar ataxias^1–10^, there are no models of most forms of ataxia. While many ataxia rodent models have been generated by genetic engineering, a small number of natural occurring ataxic mutant mice have been investigated^11–19^, contributing to understanding cerebellar function and neurodegeneration, as well as to therapeutic strategies.

We and others have reported on the *shaker* rat, a spontaneous, X-linked, recessive model of tremor and ataxia characterized by Purkinje cell (PC) degeneration^20–25^. Early work focused on characterizing the ataxia^20,26^ and PC degeneration^21^ present in the *shaker* rat, as well as developing an inbred Wistar Furth (WF) *shaker* rat strain^27^. More recent work has focused on determining the precise cause of PC loss in *shaker* rats^24,25,28^. We used the model to carry out pre-clinical studies of low-frequency deep brain stimulation of the deep cerebellum to reduce ataxia and tremor^22^, which other groups have recently continued to develop^29–31^. In these studies, we developed robust quantitative approaches for analyzing the *shaker* rat’s motor phenotype, some of which have been more recently further validated through deep neural network-based through markerless motion tracking approaches^32^.

We previously mapped the *shaker* phenotype to the end of the long arm of rat chromosome X using F2 hybrid mapping crossing *shaker* rats on the WF background with wildtype (wt) Brown Norwegian rats. This was complemented by analysis of cerebellar transcriptomes seeking to identify X-chromosomal transcripts with reduced expression, as well as coding or mis-splicing events. These studies confirmed location of the *shaker* locus on rat chromosome X using polymorphic genetic markers. This analysis narrowed the candidate gene region to 26Mbp of the telomeric region of the long arm and distal to marker DXRat21. We identified *Atp2b3* encoding PMCA3 as containing a potentially disease-coding variant. However, functional studies in yeast complementation assays did not support an effect of the amino acid substitution on function of the Ca++ pump^23^. Subsequently, we identified genetic recombination events in symptomatic F2 *shaker* rats, definitively excluding the *Atp2b3* variant as causative. We now report on additional studies combining viral, genetic, transcriptomic, and motor analysis techniques to identify the *shaker* mutation, a deletion in *Slc9a6* (solute carrier family 9, member A6) that encodes NHE6 (sodium-hydrogen exchanger 6). The deletion is predicted to lead to a truncated NHE6. Of note, *Slc9a6* is mutated in Christianson syndrome (CS), a severe epileptic encephalopathy with ataxia and autistic features^33^. We functionally confirmed the pathogenicity of the *Slc9a6* mutation by restoring NHE6 expression in PCs using viral transduction, reducing neurodegeneration and improving motor behavior.

## Results

### Brief Overview

We delineated the genetic region of interest and analyzed cerebellar and whole brain transcriptomes. We genotyped rats across the set of candidate genes and eliminated all candidate genes except *Slc9a6* through recombination events. We administered an L7-Slc9a6-GFP AAV to *shaker* rats and found that driving expression of wt NHE6 to Purkinje cells in *shaker* rats reduced cerebellar gait ataxia. We proceeded to evaluate cerebellar samples and cerebellar histology and found that administration L7-Slc9a6-GFP AAV yielded improvements in PC-specific transcripts and proteins and decreased Purkinje cell loss. Thus, we functionally demonstrated that loss-of-function mutation of *Slc9a6* was causative of the *shaker* phenotype.

### Shaker mapping

Using an F2 hybrid intercross, we mapped *shaker* to chromosome X distal to 133.45 (DxRat21) to qter. Our previous analysis of cerebellar transcriptomes revealed an amino substitution in PMCA3, encoded by *Atp2b3*, which we evaluated as a potentially disease-causing variant^23^. Although the change occurred in a highly conserved amino acid, analysis in yeast did not reveal any change in calcium transporter function. Subsequent to our publication, we observed several *shaker* WF rats that did not carry this variant. Although this clearly excluded *Atp2b3* as the causative gene, we were not able to discern whether *shaker* mapped proximal or distal to *Atp2b3*.

We developed a dual strategy to identify the *shaker* gene using genetic mapping of additional recombination events in our F2 WF/BN hybrids, as well as more effective RNA-seq analysis with greater depth and longer read lengths. To improve genetic mapping, we conducted database searches in Rnor_6.0, and Medical College of Wisconsin Rat Genome Database, and we identified 6 single-nucleotide polymorphisms in the candidate region. Of these, only rs106120845 was informative between BN and WF and mapped the *shaker* gene distal to 144.59.

To generate additional genetic markers and potentially identify the mutation itself, we used deep RNA-sequencing of cerebellar and hemispheric tissues of 3 obligate affected and 3 wt filial mates to identify coding variants, and we evaluated all variants and differentially expressed genes (DEGs). Utilizing the variant calling tool VarScan v2.3.3^34^, we found 15 changes predicted to be deleterious (3-high, 12-moderate) in 8 genes on chromosome X. Of these, 4 were frameshift variants and 11 were missense variants. Once we limited variants to the new candidate linkage region distal to 144.59, variants in three genes remained: one missense variant and splice region variant in *Flna* (filamin A), one frameshift variant in *L1cam* (L1 cell adhesion molecule), and one frameshift variant in *Slc9a6*.

All variants in the candidate genes were amplified by PCR and Sanger sequenced utilizing the WF and F2 WF/BN progeny cited in our previous publication^23^. In addition, 55 affected and 10 aged (45-week old) asymptomatic rats were screened for the *Atp2b3* and *Slc9a6* variants. The respective recombination fractions are shown in **Figure 1A**, indicating smaller recombination fractions towards the telomere. While variants in *Flna* and *L1cam* were found to have multiple recombination events, the frameshift variant in *Slc9a6* showed no recombination events in 242 shaker rats.

**Figure 1:**
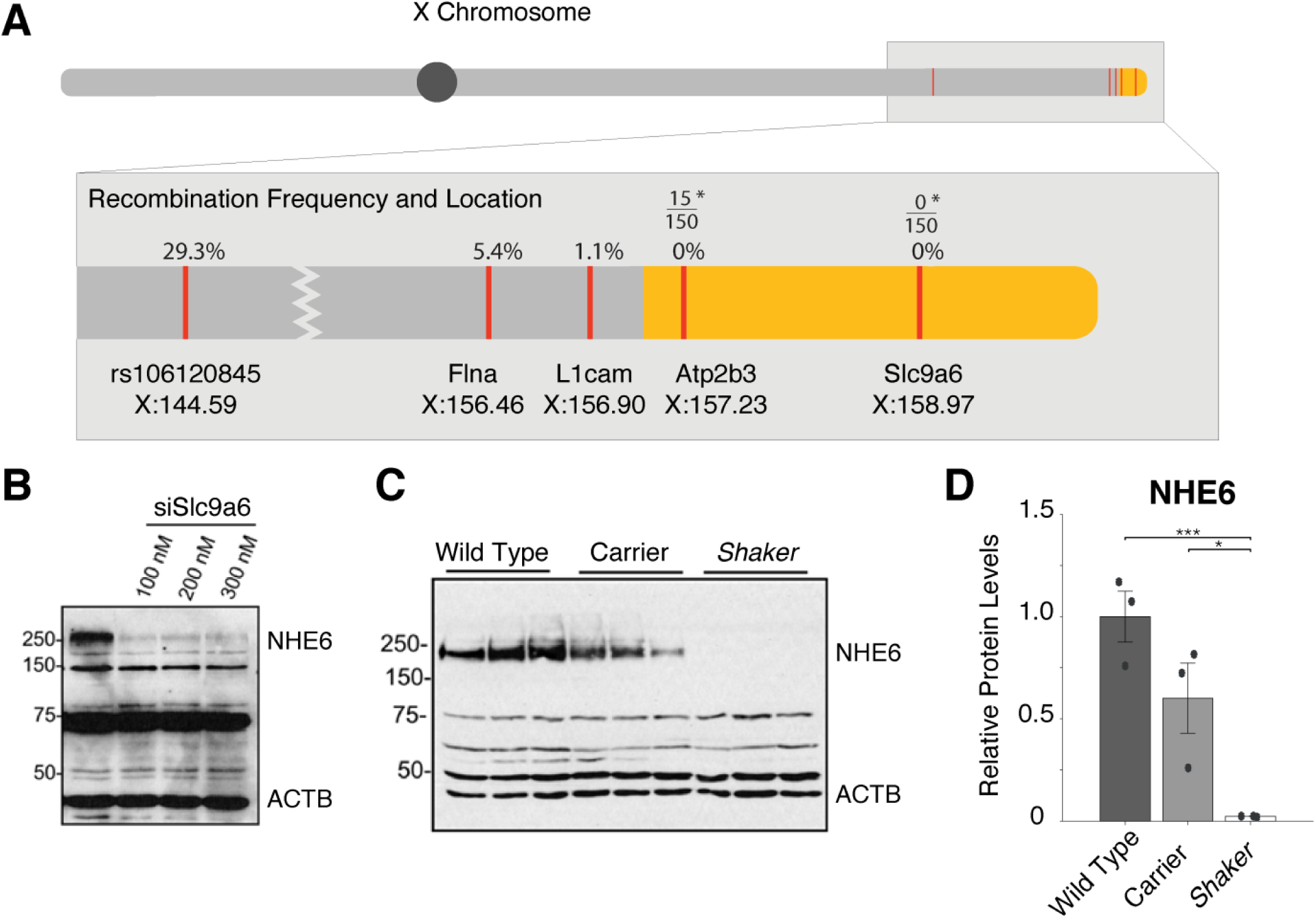
Linkage map of *shaker* and analysis of NHE6. **A**. Locations of crucial recombination events in the WF/BN F2 intercross in 92 rats are shown. Yellow refers to our narrowed region of interest. *Of 150 shaker rats in the WF background, 15 rats were wildtype for *ATP2b3*, but none were wildtype for *Slc9a6*. **B**. Western blots show that the band corresponding to glycosylated NHE6 is knocked down by siSlc9a6, validating the commercial antibody. **C.-D**. Less NHE6 expression is seen in obligate carriers than in wt; this is reduced further in *shaker* rats. Errors bars represent standard error of the mean; dots refer to individual animals.

Further confirmation for *Slc9a6* as the shaker locus came from analyzing DEGs in cerebellar transcriptomes. The top DEGs on chromosome X were *Gpr34* (G protein-coupled receptor 34), *Tlr13* (toll-like receptor 13), and *Slc9a6*. Of these top DEGs on chromosome X, *Gpr34* and *Tlr13* were well outside the linkage region, further supporting *Slc9a6* as the *shaker* allele.

### Mutation

The *Slc9a6* frameshift variant identified in *shaker* rats occurs at amino acid 64 and results in a predicted truncated protein. This frameshift was categorized as deletion-insertion variant, with base pairs 191-195 deleted, with one base pair inserted. The nomenclature is as follows: XM_217630.9(*Slc9a6*) c.[191_195delinsA], p.Ala64Glufs*23). *Slc9a6* is abundantly expressed in brain^35^, especially in PCs. In humans, mutation of human *Slc9a6* causes Christianson syndrome, a recessive X-linked epileptic encephalopathy with prominent ataxia^36^.

### NHE6 loss of function in shaker rats

After identifying *Slc9a6* as the likely *shaker* candidate gene, we tested whether the predicted NHE6 truncation would result in loss of NHE6 abundance. We evaluated several commercially available antibodies, including a rabbit polyclonal antibody that recognized several protein species at different molecular weights from 40 to 250 kDa. We sought to confirm the identity of these proteins, especially the ∼250 kDa protein, which we predicted would correspond to the fully glycosylated, dimeric form of NHE6. Indeed, while the molecular weight fluctuates with glycosylation, the fully glycosylated, dimeric form of NHE6 has been reported as 200 -250+ kDa^37,38^. We treated wells of HEK293 cells with ascending doses of *SLC9A6* siRNA^39^ and found that only the 250 kDa protein was reduced in intensity. This confirmed that the ∼250 kDa protein responded to *Slc9a6* knockdown and likely represented glycosylated NHE6 (**Figure 1B**). Of note, the 75 kDa band, which would correspond to a non-modified NHE6 moiety, did not show any change in abundance and represents a non-specific protein band, as do the other bands below 250 kDa.

To examine NHE6 expression in wt and *shaker* rats, we performed western blotting of cerebellar extracts in age matched wt, female heterozygote, and male *shaker* rats (n=3 for each at 5 weeks of age). We found gene-dosage dependent reduction of the ∼250 kDa protein in that band intensity was reduced in heterozygotes to about ∼50% and to 0% in *shaker* rats. This was consistent with loss of function caused by the frameshift *Slc9a6* mutation (**Figure 1C-D**). Notably, a lower molecular weight band representing a truncated NHE6 was not observed.

### AAV vectors for Purkinje cell targeting

To functionally test whether *Slc9a6* causes *shaker*, we developed a viral vector targeting wt rat *Slc9a6* expression to PCs and analyzed motor behavior, mRNA and protein expression, and PC morphology. We previously reported that *shaker* rats exhibited PC loss by 7 weeks of age and that PCs appear to be the predominant cell type affected early in pathogenesis^23^. Therefore, we generated a plasmid expressing a GFP-tagged rat wt *Slc9a6* under control of the PC-specific optimized mouse L7-6 (L7) promoter^40^. This plasmid was packaged in the PHP.eB capsid, ensuring widespread transduction in the CNS when injected into the lateral ventricle^41^.

To test whether ICV delivery of the L7-*Slc9a6* AAV would indeed result in expression in PCs, we generated a control vector expressing GFP under the control of the L7-6 promoter. We administered the L7-GFP AAV via ICV injection at 3 concentrations (5, 10, and 20 µL at a concentration of 2.26 × 10^11 vG/µL), as well as 10 µL of saline in 4 age-matched, wt rats. The 10 μL injection resulted in significant transduction of PCs (**Supplemental Figure 2A**), as imaged from 40 µm slices of cerebellum.

When we injected a comparable dose of Slc9a6-GFP AAV, the larger Slc9a6-GFP fusion protein did not generate detectable fluorescence, likely due to relatively low translational efficiency of the substantially larger construct and, potentially, conformational changes affecting epitope recognition. Therefore, we performed western blotting using an anti-GFP antibody. We analyzed cerebellar extracts from 4 rats administered L7-Slc9a6-GFP AAV, at 25-35 weeks of age weeks of age. While a number of non-specific bands were detected, we observed a band taht appeared to correspond to be a glycosylated NHE6 fused to GFP with a molecular weight of ≥250 kDa. This band was not observed in uninjected *shaker*, carrier, or wt rats, but was observed exclusively in *shaker* rats with motor improvement (see below). One shaker rat without motor improvement after injection also lacked a glycosylated NHE6-GFP band **Supplemental Figure 2B**).

### Motor performance is substantially improved in shaker rats treated with the L7-Slc9a6-GFP AAV

We sought functional evidence to demonstrate that mutation of *Slc9a6* was causative of the shaker phenotype. We tested whether AAV-based expression of the wt *Slc9a6* gene in *shaker* rats would prevent or reduce the motor phenotype. We injected animals with either L7-*Slc9a6-GFP AAV* or the control L7-GFP-AAV at 5 weeks of age and analyzed 30-min un-cued motor recordings taken in each rat at 7, 18, and 25 weeks of age, timepoints corresponding to the typical onset of PC loss, following the completion of PC loss, and completion of motor progression, respectively^22–24^. **Figure 2A** shows representative traces of rapid movements from rats with different genotypes and treatments at 18 weeks of age. We computed straightness of gait by taking a ratio of total distance traveled vs. displacement. At 8 weeks, there were no significant differences across groups (**Figure 2B**). At 18 weeks, *shaker* rats administered the control L7-GFP AAV were severely ataxic, while those administered L7-GFP-Slc9a6 AAV had greatly improved motor performance (**Figure 2C**). These improvements were maintained through the final time point of motor recordings, 25 weeks of age (**Figure 2D**). At all time points, uninjected wt rats were insignificantly different in gait compared to wt rats injected with L7-GFP AAV. Notably, while *shaker* rats administered the control AAV made almost exclusively uncoordinated movements at 18 and 25 weeks, several L7-Slc9a6-GFP AAV-treated *shaker* rats exhibited a strong motor therapeutic effect, being almost indistinguishable from wt rats, even at 25 weeks of age (**Figure 2E**). Representative videos taken at 18 weeks of age demonstrating the efficacy of the L7-Slc9a6-GFP AAV in improving the gait of *shaker* rats are shown in **Supplemental Video 1**.

**Figure 2:**
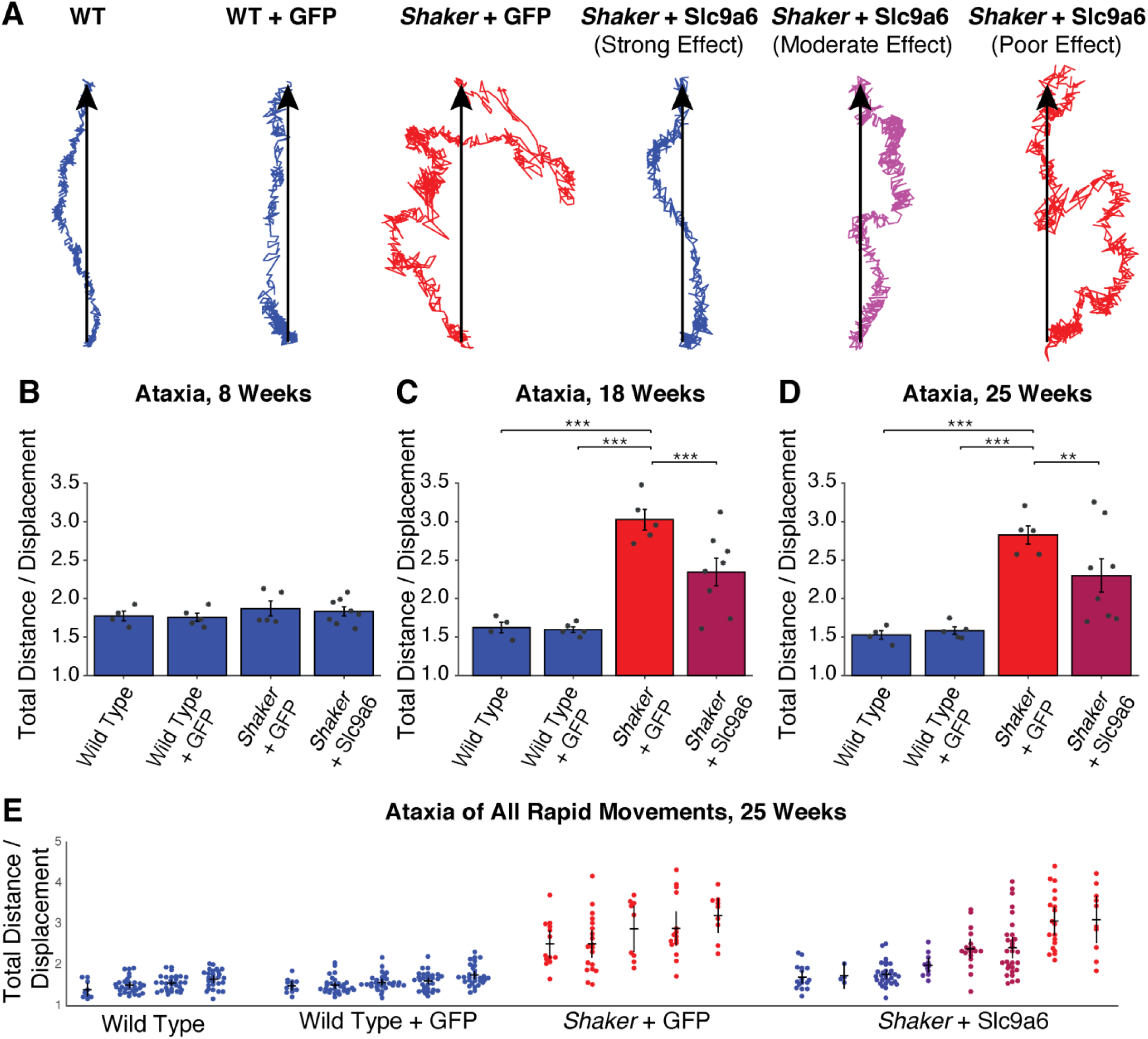
Administration of L7-Slc9a6-GFP AAV at 5 weeks of age reduces ataxic motor phenotype at later time points. **A**. Center of mass traces from representative movements made at 18 weeks of age. Treated animals are grouped based on treatment efficacy. Color scale refers to quantitative severity of ataxia. Blue refers to WT animals, red refers to untreated *shakers* (*shaker* + L7-GFP AAV). Percentage of severity from WT to untreated is represented by the shade of purple. **B-D**. Straightness of gait is computed as a measure of motor coordination. Shown is a ratio of total distance traveled vs. displacement, averaged across all rapid movements made in 30 min at each time point. Ratios are shown at **B**. 8 weeks, shortly after the typical onset of PC loss, **C**. 18 weeks, shortly after the typical completion of PC loss, and **D**. 25 weeks, at which point the motor symptom progression is typically complete. L7-GFP AAV had no impact on gait, but L7-Slc9a6-GFP AAV reduced motor dysfunction. **E**. Straightness of gait ratios are shown for all rapid movements recorded in all rats at 25 weeks. Untreated *shakers* primarily make uncoordinated movements, while treated *shakers* with a strong effect primarily make coordinated movements.

### Slc9a6 mRNA and NHE6 are partially restored in shaker rats treated with the L7-Slc9a6-GFP AAV

Cerebella from half of each control and treatment group were taken for molecular analyses. Notably, rats chosen from the treatment group for molecular analyses were representative of and spanned the group. Specifically, animals with the 1^st^, 3^rd^, 5^th^, and 8^th^ best motor treatment effect among the 8 treated animals were those analyzed at the molecular level. We performed western blotting for NHE6, identifying glycosylated NHE6 in *shaker* rats treated with L7-Slc9a6-GFP AAV (**Figure 3A&B**). Quantification of NHE6 levels (**Figure 3C**) showed significant NHE6 increases in *shaker* rats that were administered L7-Slc9a6-GFP AAV compared to those only administered L7-GFP AAV. We quantified cerebellar *Slc9a6* mRNA levels by quantitative real-time PCR (qRT-PCR) and found that compared to wt or carrier, *Slc9a6* expression was lower in *shakers* administered the control L7-GFP AAV, and this expression was partially restored by treatment with L7-Slc9a6-GFP AAV (**Figure 3D**).

**Figure 3:**
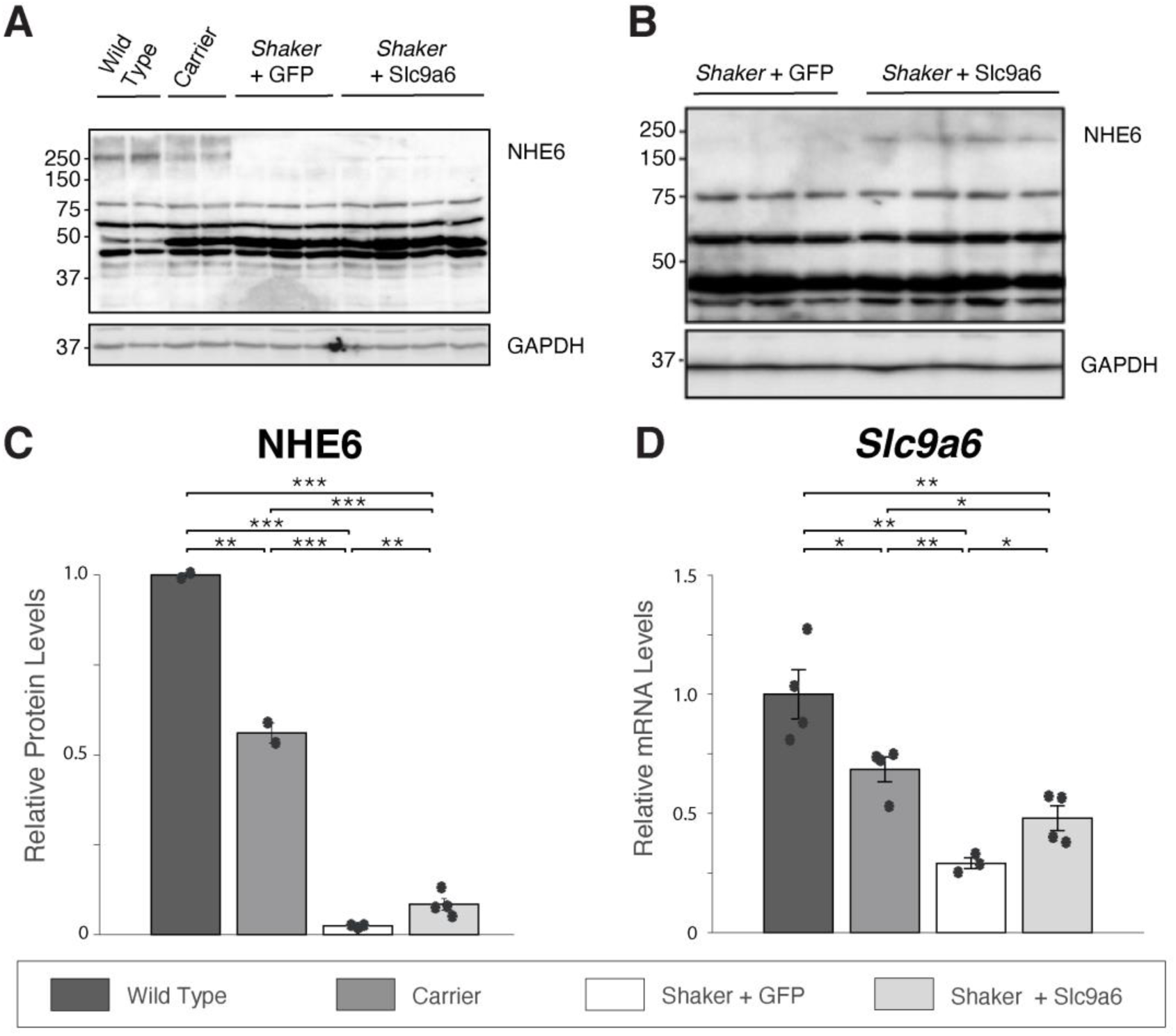
Administration of L7-Slc9a6-GFP AAV modestly, but significantly increases glycosylated NHE6 and *Slc9a6* mRNA. **A**. Western blotting of cerebellar extracts shows that the glycosylated NHE6 band is reduced in carriers, and fully reduced in *shakers* administered the control AAV. A faint band is seen in *shaker* rats treated with L7-Slc9a6-GFP AAV with ICV injection at 5 weeks of age. **B**. Repeat of **(A)** with 10x concentration of each sample confirms lack of the glycosylated NHE6 band in *shakers* +L7-GFP AAV and presence in *shakers* +L7-Slc9a6-GFP AAV. **C**. Quantified results from **(A)**. L7-Slc9a6-GFP AAV significantly restores NHE6 expression in *shaker* rats compared to those administered the control AAV. **D**. *Slc9a6* mRNA expression is evaluated through qRT-PCR. *Slc9a6* is reduced in *shaker* rats compared to wt or carrier rats, while administering L7-Slc9a6-GFP AAV increases expression.

### Purkinje cell-specific markers are partially restored in shaker rats treated with the L7-Slc9a6-GFP AAV

We quantified protein and mRNA expression for several markers associated with PC quantity and function established in prior detailed analyses of PC degeneration^3,42–47^. First, we analyzed cerebellar protein extracts by quantitative western blotting of calbindin-1 (CALB1), Regulator of G protein signaling 8 (RGS8), and Purkinje Cell Protein 2 (PCP2). We found that levels of each PC-specific protein were reduced in *shakers* compared to wt and carriers. CALB1 and RGS8 levels were partially restored by L7-Slc9a6-GFP AAV (**Figure 4A,B**), while PCP2 normalization trended towards significance (p=.062). Next, we quantified *Calb1, Rgs8*, and *Pcp2* mRNA levels by qRT-PCR. We found that all 3 mRNAs were significantly reduced in *shaker* rats compared to wt and carriers. Upon transduction with L7-Slc9a6-GFP AAV, levels were significantly increased in *shakers* (**Figure 4C**).

**Figure 4:**
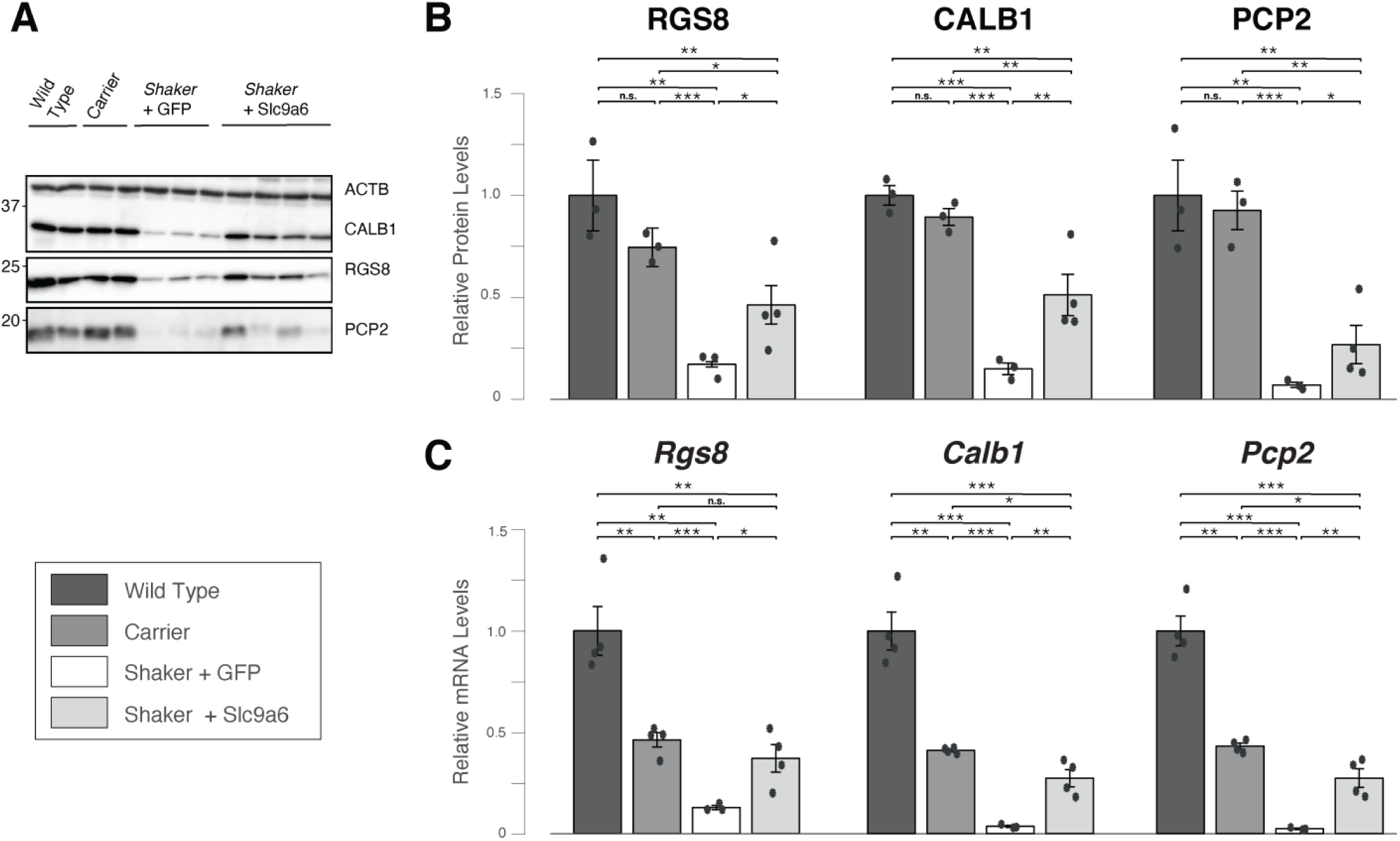
Administration of L7-Slc9a6-GFP AAV improves levels of key PC-specific mRNAs and proteins. **A**. Western blotting of cerebellar protein extracts shows reduced expression of CALB1, RGS8, and PCP2 in *shaker* rats administered the control AAV compared to wt and carriers. *Shaker* rats administered the L7-SlC9a6-GFP AAV show partially restored expression. **B**. Quantification of **(A)**. For each key PC-specific gene, administration of L7-Slc9a6-GFP AAV resulted in significantly increased expression in *shaker* rats compared to those administered the control L7-GFP AAV. **C**. mRNA expression was evaluated for *Rgs8, Calb1*, and *Pcp2* through qRT-PCR. Compared to wt, each marker was reduced significantly in carrier females and further in *shaker* rats administered the control L7-GFP AAV. *Shaker* rats administered the L7-Slc9a6-GFP AAV showed significant improvements in each marker compared to those administered the control L7-GFP AAV.

Finally, we sought to confirm these results by immunocytochemical analysis of PCs. To assess number of PC somata and extent of dendritic arborization we employed staining with an antibody to CALB1, a calcium binding protein that is highly enriched in PCs. We stained parasagittal slices from each animal sacrificed after completion of the motor experiments at 25 weeks of age with an anti-Calb1 antibody. We found that PCs in wt rats show strong CALB1 staining PC, with near total loss outside of a flocculonodular exclusion zone in *shaker* rats administered only the L7-GFP AAV. However, loss in other lobules was consistently partially prevented by treating *shaker* rats with the L7-Slc9a6-GFP AAV, matching mRNA and protein results. We show representative examples of each group in **Figure 5**.

**Figure 5:**
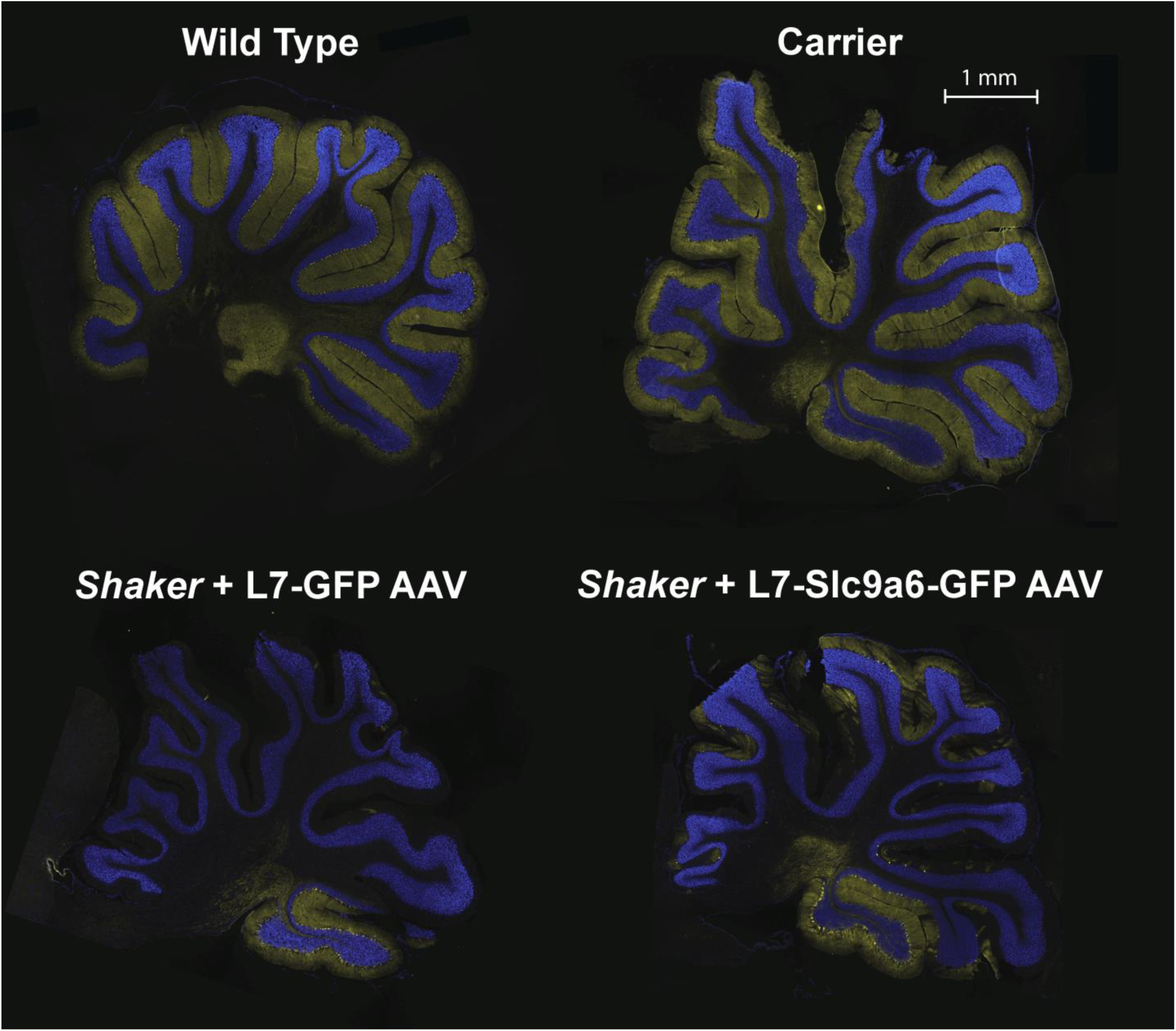
Representative 20 µm full cerebellar sagittal slices stained with Anti-Calb1 (yellow), counterstained with DAPI (blue). The L7-Slc9a6-GFP AAV prevented PC loss in *shaker* rats compared to those administered the L7-GFP control AAV. Shown are representative sagittal slices across groups. Notably, PCs tend to be preserved in the flocculonodular lobule 10. Even a relatively small degree of PC rescue led to substantial motor recovery (**Figure 2**).

## Discussion

### General

Loss-of-function mutations in endosomal Na^+^/H^+^ exchanger 6 (NHE6) encoded by *SLC9A6* cause the X-linked neurologic disorder Christianson syndrome^36^. Patients exhibit symptoms associated with both neurodevelopmental and neurodegenerative abnormalities^48^. NHE6 is mainly expressed in early and recycling endosomes and plays an important role in regulating endosomal pH^49^. Loss of NHE6 function results in overacidification of the endosome^50^, with decreased endosome-lysosome fusion in NHE6-null neurons^51^.

We identified a frameshifting mutation in *Slc9a6* that is predicted to result in an early frame shift at amino acid 64 resulting in a truncated protein of 87 amino acids. This is a spontaneous mutation that results in a tremor and ataxic phenotype, hence the designation of this strain as *shaker*. Pathologically, the phenotype is characterized by cerebellar volume loss and Purkinje cell death that proceeds from anterior to posterior, generally with less severe cell loss in a flocculonodular part of the cerebellum ^23–25^ **(Figure 5)**. We previously verified abnormal cerebellar and PC pathology in shaker rats confirming a cerebellar ataxia-like phenotype^23^.

The study objective was to identify the gene responsible for the *shaker* phenotype by linkage analysis combined with candidate gene analysis. In a previous study, we investigated *Atp2b3*, which mapped to the *shaker* candidate region. This was a promising candidate since *Atp2b3* mutations are associated with congenital cerebellar ataxia in humans^52^. However, we observed no difference in channel function by comparing wt *Atp2b3* with *Atp2b3*^R35C 23^. Additionally, recombination events observed subsequent to our publication excluded *Atp2b3*. This also refined the region to 2.7Mb, extending from 157.2 to 159.9 (Xq37-qter). While the *Slc9a6* variant did not emerge as a candidate in previous work, this current study employed both longer reads and greater depth of coverage. In addition, Slc9a6 mapped more proximal in the X chromosome in rat genomes prior to Rnor_6.0. Ultimately, based on both linkage analysis and functional evidence, we concluded that a protein-truncating mutation found in the *Slc9a6* gene is causative of the *shaker* phenotype.

### The shaker rat as a model of Christianson syndrome

At least 54 different genes on the X chromosome are linked to mental retardation^53^. One of these is *Slc9a6*, and mutations in *Slc9a6* truncating the NHE6 protein cause CS^33,36,54^, resulting in an Angelman-like, recessive, X-linked, progressive, neurodevelopmental syndrome. Symptom onset occurs in infancy, with patients exhibiting intellectual disability, autism, ataxia, lack of speech, and seizures, among other symptoms^36,48^.

Our work led to the discovery of a novel PTV (p.A64Efs*23) in *Slc9a6* in the *shaker* rat, making this the first rat model of CS. Other groups have worked with *Slc9a6* knockout mice as well as *Slc9a6* mutant mice^37,55,56^ to demonstrate that *Slc9a6* loss of function causes the relevant defects, including disrupted endosomal-lysosomal function, neurodevelopmental and neurodegenerative pathology, impaired plasticity, and impoverished neuronal arborization^37,37,50,55,57–60^. Further, *Slc9a6* haploinsufficient mice develop PC, motor, and visuospatial abnormalities^61^. Other *Slc9a6* rodent models are not reported as recapitulating all components of the human disease, such as seizures, cognitive deficits, or autistic behavior. Given that rats have a complex social repertoire, the *shaker* rat may be well suited for detailed dissection of the behavioral / cognitive components of CS.

### NHE6 function in the context of the shaker rat

NHE6 is a membrane-bound protein functioning as a Na^+^/H^+^ exchanger, providing a leak pathway for protons and limiting endosomal acidification.^62^ NHE6 expression is high in PCs and coincides with PC loss in NHE6-null mice.^59^ NHE6 mutation leads to neuronal degeneration through apoptosis^37^, which is the mechanism for PCs death in the *shaker* rat^24,25^. However, NHE6 expression is substantially more widespread than just PCs, and loss of function could potentially impact the entire nervous system.^58^ Thus, future work in the *shaker* rat should include an evaluation of NHE6 deficits in cerebrum and spinal cord.

### Functional evidence suggesting that Slc9a6 mutation underlies the shaker phenotype

In this work, we generated an AAV vector targeting expression of *Slc9a6* to PCs using the L7 (Pcp2) promoter. AAV administration after P1-P2 in rodents can result in relatively low neuronal yield^63^. Indeed, our ICV injection at 5 weeks of age provided relatively low levels of glycosylated NHE6 compared with levels in wt rats, ∼10% of wt cerebellar NHE6 (**Figure 3**). However, despite relatively low cerebellar NHE6, PC-specific proteins and mRNA levels, markers of PC health, improved substantially toward normalization (**Figure 4**). It is interesting that key mRNAs and proteins decreased in SCA2 mouse models are also decreased in *shaker*. Although it is known that these changes occur prior to PC cell loss or significant rarification of PC dendrites in SCA2 mouse models^42^, future studies will be needed to examine transcriptomic changes across different time points.

Behavioral analyses that we performed used a previously described force plate paradigm that allows analysis of tremor and gait ataxia at a spatial and temporal resolution exceeding that of video-based analysis^22^. Ataxic phenotypes significantly improved, with a subset of rats approaching near-normal motor function (**Figure 2**). These functional data strongly support *Slc9a6* as the gene responsible for the shaker phenotype.

### The prospect of gene therapy for Christianson syndrome

Viral gene therapy is an attractive strategy for treating CS, supported by this study. However, therapeutic optimization of the viral construct will be required. First, a therapeutic construct would lack a GFP tag; given the large size of our construct with the GFP tag, it is likely that translational efficiency would be substantially better in this proposed construct, which would likely yield greater therapeutic benefit. Further, using a promoter driving expression beyond just Purkinje cells is almost certainly necessary to treat a number of other CS symptoms. With this in mind, using a pan-neuronal promoter such as CAG may be ideal^64^. However, NHE6 gain of function has been reported as over-alkalizing endosomes, yielding neuronal atrophy^65^. Further, off target effects may occur, particularly with improper dosing. Therefore, precise dose response studies will be of high importance for therapeutic development.

### NHE6 in the context of Alzheimer’s disease

The Apolipoprotein ε4 (ApoE4) variant is the greatest genetic risk factor for Alzheimer’s disease^66–68^. Several recent studies have focused on the interactions of NHE6 and ApoE4, and it is clear that NHE6 has a major role in ApoE4-mediated neurodegeneration. Recent work has shown that NHE6 knockout mice exhibit elevated amyloid beta. Endosomal acidification in ApoE4 astrocytes is caused by NHE6 downregulation, leading to reduced amyloid beta clearance, suggesting NHE6 overexpression as a potential therapy for AD.^57^ Further, NHE6 expression attenuated amyloid precursor protein processing and amyloid beta secretion in a HEK293 cell model of human amyloid precursor protein expression^62^. In contrast, conditionally depleting NHE6 *in vivo* post-developmentally corrected ApoE4-mediated synaptic impairments and prevented amyloid beta production in mice^69^. Thus, both decreased^69^ and increased^57^ NHE6 expression have been proposed as potentially therapeutic for Alzheimer’s disease. ApoE4 also exacerbates tauopathies^70^, and CS is tied to 4R tauopathy^71^. Thus, NHE6 may play a role in both amyloid and tau pathogenesis. Additional studies will be necessary to fully elucidate how NHE6 might be therapeutically targeted, giving the *shaker* rat additional investigative significance well beyond CS.

### Phenotypic variation across the shaker rat, NHE6-null mice, and humans with Christianson syndrome

CS patients universally have profound cognitive deficits. While we do not report on cognitive testing here, we have not qualitatively noted behavioral abnormalities of a similar scale in *shaker* rats. Similarly, *NHE6* knockout mice have not been reported as replicating the severity of cognitive deficits observed in CS. In terms of PC degeneration, reports of NHE6-null mice indicate a slower progression, perhaps with less severe PC loss.^59,72^ Among the possible underlying mechanisms, one involves differential activity of NHE6 paralogs such as NHE9, which is also localized in recycling endosomes. Indeed, there exists a complex interplay between vesicular ATP-dependent proton pumps and the NHEs, and it is possible that any number of modifiers may yield important differences in endosomal pH and PC health.

### Potential Caveats

NHE6 protein levels in treated *shaker* rats significantly increased but remained relatively low. We interpret this as being related to restored NHE6 expression only in Purkinje cells, but not astrocytes. Perhaps the use of a ubiquitous promoter would restore NHE6 closer to wt levels, but it remains clear that even some degree of restoration is sufficient for substantial protection of motor function.

Successful expression of the L7-Slc9a6-GFP AAV was determined through western blotting rather than histology. While the L7-GFP AAV yielded strong histological signal (**Supplemental Figure 2A**), GFP expressed by the L7-Slc9a6-GFP AAV was not clearly detectable over background autofluorescence by fluorescent microscopy. This may have been due to either low expression or misfolding of the fused GFP. Importantly, however, a specific band corresponding to the Slc9a6-GFP fusion protein was detected upon western blotting the cerebella of L7-Slc9a6-GFP AAV-treated *shaker* rats (**Supplemental Figure 2B**) and was exclusive to rats with therapeutic benefit. Further, expression was also validated through *Slc9a6* mRNA and NHE6 protein expression, both of which were increased significantly in cerebellum. This yielded significant changes in numerous key PC proteins associated with very substantial improvement in motor performance and indicating that the L7-6 targeting remained intact in L7-Slc9a6-GFP. As we were approaching AAV packaging limits with our construct, this likely substantially impacted translational efficiency.^73^ Future work using the L7 promoter would benefit from a dual virus injection consisting of primarily L7-Slc9a6 AAV and a small amount of L7-GFP AAV to validate viral presence in cerebellum. This approach would enable both visualization of transduction and greater translational efficiency.

### Conclusions

The cerebellar dysfunction and motor symptoms exhibited by the *shaker* rat are caused by loss-of-function mutation in the *Slc9a6* gene. Thus, the *shaker* rat is a spontaneous model of CS. Given the *shaker* rat’s strong motor phenotype, it should be considered as a model for mechanistic study and therapeutic development for CS.

## Materials and Methods

All animal work was done under an approved IACUC protocol at the University of Utah. Rats were housed 2-5 per cage under a 12:12 h light cycle with food and water provided *ad libitum*. We maintain the *shaker* colony in the Wistar Furth (WF) inbred background (Envigo RMS LLC).

### Genetic mapping

The *shaker* rat used in these studies is a spontaneous X-linked recessive mutant that we have previously described^23^. Once the previously identified candidate gene *Atp2b3* was excluded, we refined to 2.7Mb, extending from 157.2 to 159.9 (Xq37-qter) through an F2, WF / Brown Norwegian (BN) intercross. The F1 offspring were generated by breeding affected WF males and commercial wt BN female rats, and the F2 generation was obtained by breeding affected F1 males with obligate carrier females. An assumption was made that the WF alleles would segregate in close proximity to the causative *shaker* gene. An additional 55 WF affected males and ten 45-week old asymptomatic (non *Slc9a6* variant positive) N6F0 WF were added to our previously reported cohort for a total 242 affected male rats: 150 WF and 92 F2 WF/BN.

### RNA-Seq

We conducted deep RNA-seq using total RNA on whole cerebella and brain hemispheres of 3 wt and 3 obligate affected 5-week old WF males. We extracted total RNA with miRNAeasy minikit (Qiagen Inc., Valencia, CA), obtaining RNA quality using the Agilent RNA ScreenTape Assay (Agilent; Santa Clara, CA). Library preparation took place utilizing Illumina TruSeq Stranded RNA Kit with Ribo-Zero Gold (rat). We generated paired-end 125-bp reads on a Hiseq 2000 machine at the University of Utah Microarray and Genomic Analysis Shared Resource using Illumina HiSeq 125 cycle paired-end sequencing v4.

We used SAMTools Mpileup on the alignment files and VarScan v2.3.3^34^ for variant calling. Per our exclusion of *Atp2b3* as a causative disease gene, we focused on variants found on ChrX: 157,239,000 to qter. In order to estimate variance-mean dependence and test for differential expression, we utilized DESeq2.^74^

We examined the relative expression of X chromosome genes to look for differential expression and logarithmic fold changes. We first sorted our data by Max DESeq2 AdjP values and then by then by Log2 Ratio. The Max DESeq2 AdjP values were then defined as the maximum observed -10Log10(AdjP) between any pairwise DESeq2 comparisons. All p-values and AdjP were phred transformed where the value is - 10*Log10(AdjP or pval); thus -10*Log10(0.05) = 13. Therefore, significant cutoffs are those of values equal to or greater than 13. The Log2 ratio is defined as log_2_ (T/R), where T is the gene expression level in affected rats and R is the gene expression level in the wt samples.

### Mutation Description

To generate standardized nomenclature for this protein truncating variant (PTV), we utilized the freely available Mutalyzer-Description extractor software to generate the Human Genome Variation Society (HGVS) variant description. Although the NCBI Reference sequence XM_217630.9 is more reflective of the *Slc9a6* human transcript, the most recent Rat Genome release Rnor_6.0 has a transcript different at the 5’ end of the observed mutation adding an additional 20 amino acids (ENSRNOT00000066809.3) (**Supplemental Figure 1**). Because Mutalyzer-Description extractor software is only compatible with NCBI sequences, we used NCBI to generate the variant description and Rnor_6.0 to anchor the markers used for genetic mapping.

### Genotyping Methods

We isolated genomic DNA from rat tail tips using Qiagen genomic DNA extraction kit (Qiagen Inc., USA). We performed genotyping via PCR; a 363 bp fragment was produced with the following primers: forward of 5’-AAGACATGGCTGTGGCTCGG-3’ and reverse of 5’-AGCTAGGGGACAGGGGTCCG-3’. Due to the minimal difference in fragment sizes (4bp), we confirmed genotype via Sanger sequencing with the same forward primer.

### AAV Production

We derived rat cDNA sequences for *Slc9a6* (XM_001053956) from the NCBI DNA database, which we used to design primers to PCR-amplify the coding sequences from a cDNA library made from wt rat cerebellar RNA. The primer set was as follows: Slc9a6-F: 5’-TTTATGGCTGTG GCTCGGCGCGGCTGG-3’ and Slc9a6-R: 5’-TTTCGGCTGGACTGTGTCTTGTGTCATC-3’. We cloned the amplified PCR product into TOPO vector (TOPO™ TA Cloning™ Kit, ThermoFisher, Cat# K457502) and verified by sequencing. The adeno associated virus, pAAV/L7-6-GFP-WPRE (woodchuck hepatitis post transcriptional regulatory element)-SV40, was a gift from Dr. Hirokazu Hirai^40^, and the pUCmini-iCAP-PHP.eB was designed by the Gradinaru group (Addgene, plasmid #103005)^41^. For the PHP.eB AAV expression plasmid, the *Slc9a6* cDNA was re-PCR amplified from TOPO cloning vector using *Age*I restriction sites and subsequently cloned into pAAV/L7-6-GFP-WPRE plasmid at *Age*I site, designated as pAAV/L7-6-Slc9a6-GFP. All constructs were verified by sequencing. We generated recombinant AAV particles in the Drug Discovery Core Facility, University of Utah. Briefly, HEK-293T cells were co-transfected with the following plasmids: pAAV/L7-6-Slc9A6-GFP or pAAV/L7-6-GFP (control) with pHelper (Stratagene, La Jolla, CA, USA), and pUCmini-iCAP-PHP.eB followed by viral particles purification, concentration and genomic titer determination.

### Surgical Procedures

18 age-matched rats were administered either the L7-GFP control AAV or the L7-Slc9a6-GFP AAV, including 3 male wt rats, 2 female carriers, 6 males hemizygous, and 7 females homozygous for the candidate allele. We administered *the* L7-Slc9a6-GFP AAV to 8 *shakers*, and the control L7-GFP AAV to 5 *shakers* and 5 wt or carriers. An additional 4 rats – 2 male wt and 2 female carriers – served as uninjected controls. Littermate effect was avoided by group randomization. The wt rats were all male given that our focus was on treatment of *shaker* rats and the fact that it is not possible to generate wt and *shaker* females in the same litter. Notably, we previously analyzed the gait of wt males and females and found no significant differences in straightness of gait between wt animals of both sexes^22^. We anesthetized rats with 2% isoflurane at 35 +/- 2 days of age. We shaved and disinfected the surgical site and placed rats on an electronically controlled heating pad set to body temperature in a stereotactic frame. We administered 0.1 mL bupivacaine to the incision site and opened to the scalp and dried. We marked the craniotomy site for the injection of AAV to the right lateral ventricle with respect to bregma and used a burr to drill a hole in the marked location. We inserted a needle attached to a syringe (Hamilton Co., Reno, NV) into the lateral ventricle, waited for 3 minutes, then injected 10 µL of virus at < 2.0 µL / min. Five minutes after injection, the needle was withdrawn at a rate not exceeding 1.0 mm / min. We sutured the incision and provided 0.1 mg/kg carprofen subcutaneously daily for 3 days. Rats were then given a 12- to 16-day recovery period.

### Motor Analyses

We made 30-minute force plate-based motor recordings in the aforementioned rats at 7, 18, and 25 weeks of age using previously described methods^22,76^. These timepoints chosen were based on the typical start of PC degeneration, after the end of the typical PC degeneration, and when motor symptom progression is complete^22^. We tracked center of mass at 1000 Hz and analyzed straightness of gait during rapid movements as a measure of ataxia of gait. We computed a ratio of distance traveled to displacement during rapid movements, in which we previously showed that wt rats average 1.5-1.7 and severely ataxic rats average ∼3.0^22^. Notably, we previously found that carrier rats have insignificantly different motor ataxia from wt rats. It is impossible to breed both affected females and wt females in the same litter. Thus, in the interest of litter matching across groups, motor results labeled as wt are a combination of wt and carrier.

### Extraction of tissues for cellular and molecular analyses

Following experiments, we deeply anesthetized rats and extracted all cerebella. In 21 of 22 rats used in AAV experiments, we retained one of half of each brains for histological analyses, while one brain was not evaluated due to damage on removal. We took the other half of the 11 of the 22 cerebella – as well as 2 more uninjected, age-matched wt and carrier rats – for a total of 15 samples and flash froze at -80 °C. These samples were homogenized, with half of each sample used for mRNA analyses and the other half used for protein analyses.

### Preparation of protein lysates and western blotting

We prepared cellular extracts by a single-step lysis method.^75^ We suspended harvested cells in SDS-PAGE sample buffer [Laemmli sample buffer (Bio-Rad, Cat# 161-0737)] before boiling for 5 min. We used equal amounts of the extracts for western blot analyses. We prepared rat cerebellar protein extracts by homogenization of tissues in extraction buffer [25 mM Tris-HCl pH 7.6, 300 mM NaCl, 0.5% Nonidet P-40, 2 mM EDTA, 2 mM MgCl_2_, 0.5 M urea, and protease inhibitors (Sigma-Aldrich, P-8340)], followed by centrifugation at 4°C for 20 min at 14,000 RPM, only using supernatants for western blotting. We resolved protein extracts by SDS-PAGE and transferred to Hybond P membranes (Amersham Bioscience) before processing for western blotting according to our previously published protocol.^75^ We used the Immobilon Western Chemiluminescent HRP Substrate (EMD Millipore, Cat# WBKLSO500) to visualize the signals and detected on the ChemiDoc MP imager (Bio-Rad). For some blots, we used a film developing system and quantified band intensities using ImageJ software after inversion of the images. Relative protein abundances were expressed as ratios to ACTB or GAPDH.

We used the following antibodies and dilutions for western blotting: Anti-NHE-6/*Slc9a6* antibody [(1: 5000), Abcam, Cat# ab137185), GFP antibody (B-2) [(1:2,000), Santa Cruz, sc-9996], GAPDH (14C10) rabbit mAb [(1:5,000), Cell Signaling, Cat# 2118], monoclonal anti-β-Actin−peroxidase antibody (clone AC-15) [(1:30,000), Sigma-Aldrich, A3854], monoclonal Anti-Calbindin-D-28K antibody [(1:5,000), Sigma-Aldrich, C9848], RGS8 antibody [(1:5,000) (Novus Biologicals, NBP2-20153)], PCP-2 antibody (F-3) [(1:3,000), Santa Cruz, sc-137064], and GFP rabbit polyclonal antibody (FL) [(1:1,000), Santa Cruz, sc-8334]. The secondary antibodies were peroxidase-conjugated AffiniPure goat anti-rabbit IgG (H + L) antibody [(1:5000), Jackson ImmunoResearch Laboratories, Cat# 111-035-144] and Peroxidase-conjugated horse anti-mouse IgG (H+L) antibody [(1:5,000) (Vector laboratories, PI-2000)]. Initial western blots were carried out using 3 wt rats of both genders, 3 carrier females, and 3 *shaker* rats of both genders, while later western blots were made from the rats described below, as well as 2 additional uninjected, age-matched wt rats, 1 male and 1 female, and 2 additional carrier females.

### *siRNAs* and reagents

For siRNA experiments, we used the following siRNAs: All Star Negative Control siRNA (Qiagen, Cat# 1027280), human *siSLC9A6:* 5’-GGA ACA GCA AUU UCU UGU UUC GUUA-3’ ^39^. siRNA oligonucleotides were synthesized by Invitrogen, USA. The oligonucleotides were deprotected and the complementary strands were annealed.

### RNA Analyses

Total RNA from flash frozen cerebellums was extracted using the RNeasy Mini Kit (Qiagen) according to the manufacturer’s protocol. The extracted RNA was then used to synthesize cDNA using ProtoScript® First Strand cDNA Synthesis Kit (New England Biolabs). Quantitative real-time PCR (qRT-PCR) was performed in an Applied Biosystems™ QuantStudio™ 12K Flex Real-time PCR system using TaqMan™ Gene Expression Assays (FAM) (ThermoFisher, actin Rn00667869_m1, slc9a6 Rn01402034_m1, Calb1 Rn00583140_m1, fam107b Rn0165005_m1, pcp2 Rn01403231_g1, rgs8 Rn00571066_m1, gfap Rn01253033_m1) and TaqMan™ Fast Advanced Master Mix (ThermoFisher). PCR cycling parameters were set according to the manufacturer’s protocol for TaqMan™ Fast Advanced Master Mix and TaqMan™ Gene Expression Assays. The qPCR data was analyzed by the ΔΔCt method, using ΔΔCt values calculated by the QuantStudio™ 12K Flex software.

### Histological Analyses

We incubated half of each cerebellum in 4% PFA for 24 hours followed by 24 hours at each of 10%, 20%, and 30% sucrose optimal cutting temperature (OCT) blocks were cut into 20 μm coronal slices with a Leica 5100S cryostat (Wetzlar, Germany), retaining 1 slice every 200 μm. In all 21 animals, we stained at least 3 slices with a mouse monoclonal anti-calbindin-D-28K (1:500) and a counterstain for DAPI.

### Statistical Analyses

Comparisons were made via 2-sample Student’s t-tests with n = the number of animals unless otherwise noted. P-values are reported via significance bars for all mentioned comparisons as follows: ns refers to p>0.05, while * refers to p<0.05, ** refers to p<0.01, and *** refers to p<0.001.

## Acknowledgements

This work was supported by NIH NINDS R37NS033123 (Pulst), NIH NINDS R21NS104799 (Pulst), NIH NINDS R21NS079852 (Pulst), an RTW Charitable Foundation Grant (Pulst, Anderson, Paul), a National Ataxia Foundation Postdoctoral Fellowship (Anderson), a National Ataxia Foundation Junior Investigator Award (Anderson), and a Harrington Discovery Institute Rare Disease Scholar Award (Scoles). The authors thank Christopher R. Nelson for his technical assistance with histological slice preparation. We acknowledge the Cell Imaging Core at the University of Utah for use of several microscopes and thank Dr. Michael Bridge for his assistance.

---

**Supplemental Figure 1:**
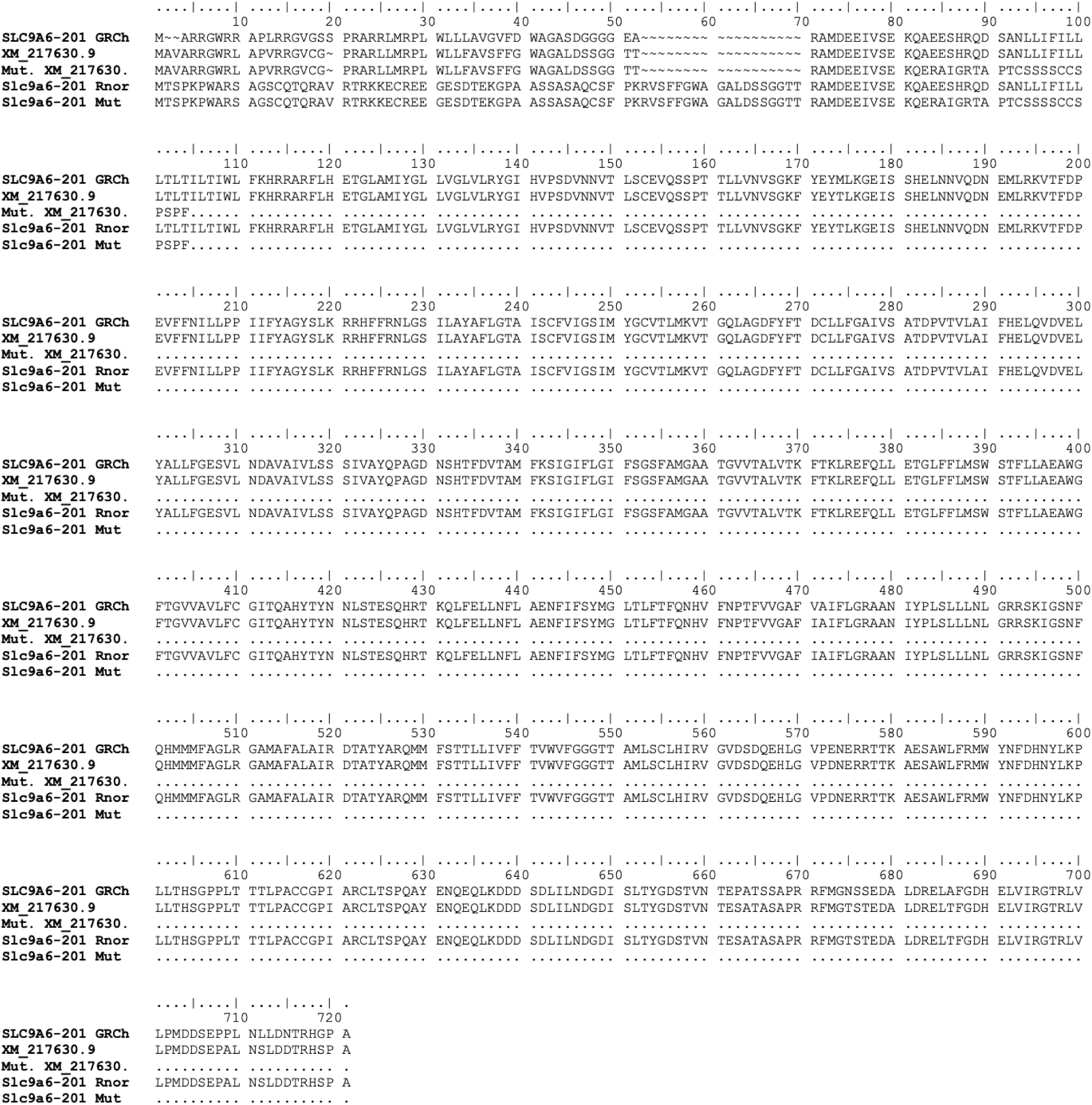
Human and rat amino acid alignment. Shown in order from top to bottom are the protein sequence of the *SLC9A6* Isoform X1 (SLC9A6-201), the NCBI *Slc9a6* rat protein sequence (XM_217630.9), the predicted mutated NCBI rat aa sequence, the rat genome release Rnor_6.0 *Slc9a6* rat protein sequence, and its predicted mutated sequence.

**Supplemental Figure 2:**
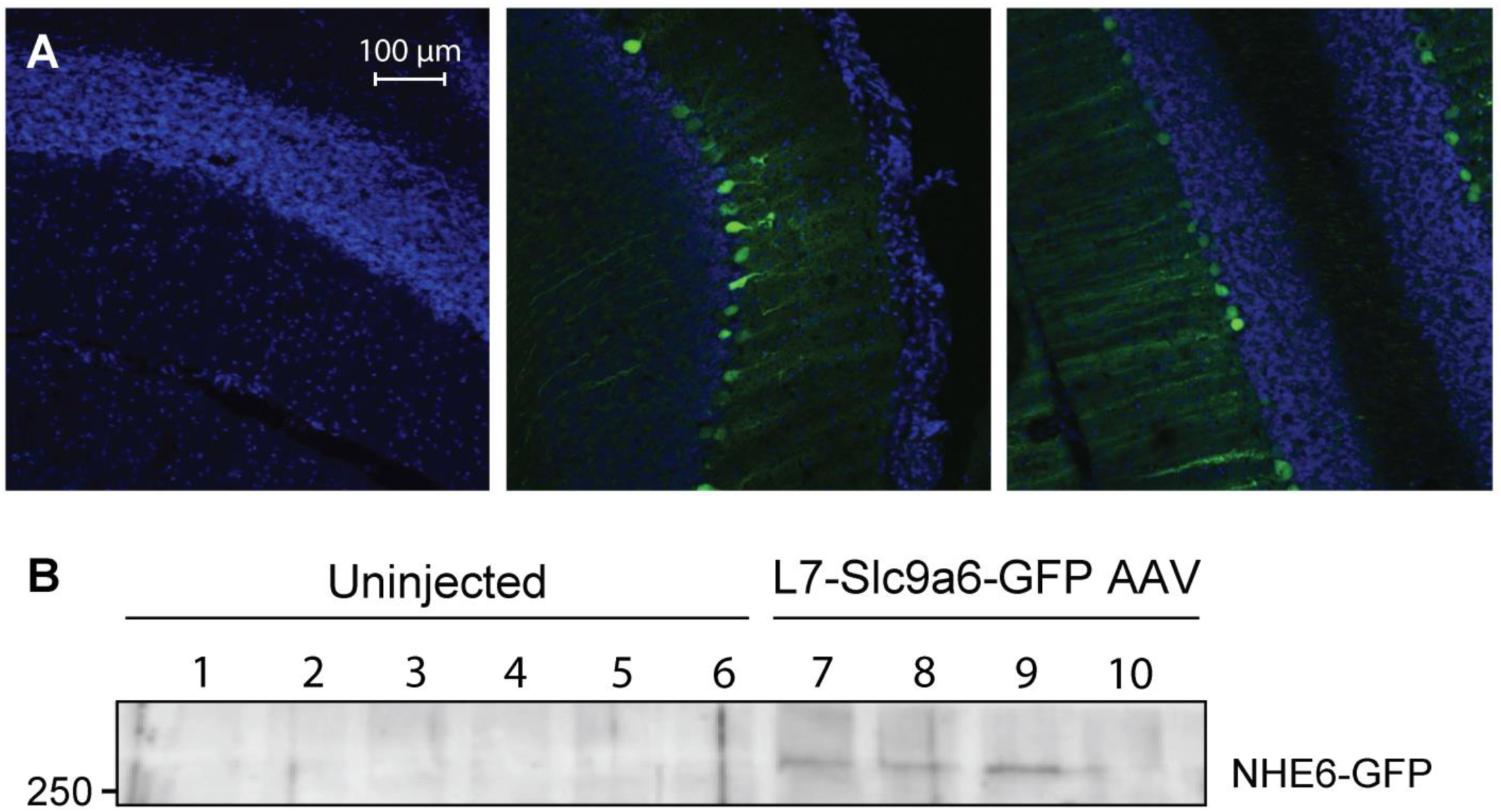
Transduction of PCs. **A**. We verified target engagement of L7-GFP AAV via microscopy. Left: 10 µL saline administered via ICV injection; Middle: 10 µL L7-GFP; Right: 20 µL L7-GFP. **B**. A specific signal was not seen after L7-Slc9a6-GFP administration, whether due to low translational efficiency or conformational changes in the fused NHE6-GFP. Therefore, we evaluated transduction by western blotting, using a rabbit polyclonal anti-GFP antibody. Shown here are 6 uninjected wt rats (lanes 1-6) and 4 *shaker* rats administered the L7-Slc9a6-GFP AAV (lanes 7-10). Lanes 7-9 showed a faint, but specific signal at 260-270 kDa, consistent with the expected size of glycosylated NHE6 fused to GFP, while lane 10 did not. Notably, the sample from lane 10 was taken from an animal with no detectable therapeutic benefit in terms of motor performance (8^th^ highest ataxia score in 8 animals administered L7-Slc9a6-GFP AAV, **Figure 2**), with unchanged RGS8 and PCP2 levels (rightmost lane, **Figure 4A**). This indicates a strong likelihood that the AAV was not properly administered in this animal.

<u>Supplemental Video 1:</u> Rats are shown at 18 weeks of age. We show 3 untreated *shaker* rats, i.e. administered only the L7-GFP AAV, 2 *shaker* rats quantitatively determined to have ∼90% reduction of motor symptoms compared to untreated *shaker* rats, and 2 treated *shaker* rats – i.e. administered the L7-Slc9a6-GFP AAV with median treatment efficacy. Obvious differences across the groups are apparent.

## References

1. Burright, E. N. et al. SCA1 transgenic mice: A model for neurodegeneration caused by an expanded CAG trinucleotide repeat. Cell 82, 937–948 (1995).

2. Huynh, D. P., Figueroa, K., Hoang, N. & Pulst, S.-M. Nuclear localization or inclusion body formation of ataxin-2 are not necessary for SCA2 pathogenesis in mouse or human. Nat. Genet. 26, 44–50 (2000).

3. Dansithong, W. et al. Ataxin-2 Regulates RGS8 Translation in a New BAC-SCA2 Transgenic Mouse Model. PLoS Genet. 11, (2015).

4. Colomer Gould, V. F. Mouse Models of Spinocerebellar Ataxia Type 3 (Machado-Joseph Disease). Neurotherapeutics 9, 285–296 (2012).

5. Watase, K. et al. Spinocerebellar ataxia type 6 knockin mice develop a progressive neuronal dysfunction with age-dependent accumulation of mutant CaV2.1 channels. Proc. Natl. Acad. Sci. 105, 11987–11992 (2008).

6. Yoo, S. Y. et al. SCA7 knockin mice model human SCA7 and reveal gradual accumulation of mutant ataxin-7 in neurons and abnormalities in short-term plasticity. Neuron 37, 383–401 (2003).

7. White, M. et al. Transgenic mice with SCA10 pentanucleotide repeats show motor phenotype and susceptibility to seizure — A toxic RNA gain-of-function model. J. Neurosci. Res. 90, 706–714 (2012).

8. Cendelin, J. et al. Consensus Paper: Strengths and Weaknesses of Animal Models of Spinocerebellar Ataxias and Their Clinical Implications. The Cerebellum (2021) doi:10.1007/s12311-021-01311-1.

9. Ramani, B. et al. A knockin mouse model of spinocerebellar ataxia type 3 exhibits prominent aggregate pathology and aberrant splicing of the disease gene transcript. Hum. Mol. Genet. 24, 1211–1224 (2015).

10. Ramani, B. et al. Comparison of spinocerebellar ataxia type 3 mouse models identifies early gain-of-function, cell-autonomous transcriptional changes in oligodendrocytes. Hum. Mol. Genet. 26, 3362–3374 (2017).

11. Phillips, R. J. S. ‘Lurcher’, a new gene in linkage group XI of the house mouse. J. Genet. 57, 35 (1960).

12. Lalouette, A., Guénet, J. L. & Vriz, S. Hotfoot mouse mutations affect the delta 2 glutamate receptor gene and are allelic to lurcher. Genomics 50, 9–13 (1998).

13. Sidman, R. L., Lane, P. W. & Dickie, M. M. Staggerer, a new mutation in the mouse affecting the cerebellum. Science 137, 610–612 (1962).

14. Hamilton, B. A. et al. Disruption of the nuclear hormone receptor RORalpha in staggerer mice. Nature 379, 736–739 (1996).

15. Falconer, D. S. Two new mutants, ‘trembler’ and ‘reeler’, with neurological actions in the house mouse (Mus musculus L.). J. Genet. 50, 192–201 (1951).

16. Patil, N. et al. A potassium channel mutation in weaver mice implicates membrane excitability in granule cell differentiation. Nat. Genet. 11, 126–129 (1995).

17. Sweet, H. O., Bronson, R. T., Johnson, K. R., Cook, S. A. & Davisson, M. T. Scrambler, a new neurological mutation of the mouse with abnormalities of neuronal migration. Mamm. Genome Off. J. Int. Mamm. Genome Soc. 7, 798–802 (1996).

18. Cendelin, J. From mice to men: lessons from mutant ataxic mice. Cerebellum Ataxias 1, (2014).

19. Manto, M. & Marmolino, D. Animal Models of Human Cerebellar Ataxias: a Cornerstone for the Therapies of the Twenty-First Century. The Cerebellum 8, 137–154 (2009).

20. La Regina, M. C., Yates-Siilata, K., Woods, L. & Tolbert, D. Preliminary characterization of hereditary cerebellar ataxia in rats. Lab. Anim. Sci. 42, 19–26 (1992).

21. Tolbert, D. L., Ewald, M., Gutting, J. & La Regina, M. C. Spatial and temporal pattern of Purkinje cell degeneration in shaker mutant rats with hereditary cerebellar ataxia. J. Comp. Neurol. 355, 490–507 (1995).

22. Anderson, C. J., Figueroa, K. P., Dorval, A. D. & Pulst, S. M. Deep cerebellar stimulation reduces ataxic motor symptoms in the shaker rat. Ann. Neurol. (2019) doi:10.1002/ana.25464.

23. Figueroa, K. P. et al. Spontaneous shaker rat mutant – a new model for X-linked tremor/ataxia. Dis. Model. Mech. 9, 553–562 (2016).

24. Erekat, N. S. Cerebellar Purkinje cells die by apoptosis in the shaker mutant rat. Brain Res. 1657, 323–332 (2017).

25. Erekat, N. S. Autophagy precedes apoptosis among at risk cerebellar Purkinje cells in the shaker mutant rat: an ultrastructural study. Ultrastruct. Pathol. 42, 162–169 (2018).

26. Wolf, L. W., LaRegina, M. C. & Tolbert, D. L. A behavioral study of the development of hereditary cerebellar ataxia in the shaker rat mutant. Behav. Brain Res. 75, 67–81 (1996).

27. Clark, B. R., LaRegina, M. & Tolbert, D. L. X-linked transmission of the shaker mutation in rats with hereditary Purkinje cell degeneration and ataxia. Brain Res. 858, 264–273 (2000).

28. Erekat, N. S. Active caspase-3 upregulation is augmented in at-risk cerebellar Purkinje cells following inferior olive chemoablation in the shaker mutant rat: an immunofluorescence study. Neurol. Res. 41, 234– 241 (2019).

29. Miterko, L. N. et al. Neuromodulation of the cerebellum rescues movement in a mouse model of ataxia. Nat. Commun. 12, 1295 (2021).

30. Cury, R. G. et al. Effects of dentate nucleus stimulation in spinocerebellar ataxia type 3. Parkinsonism Relat. Disord. 69, 91–93 (2019).

31. Cury, R. G. et al. Safety and Outcomes of Dentate Nucleus Deep Brain Stimulation for Cerebellar Ataxia. The Cerebellum (2021) doi:10.1007/s12311-021-01326-8.

32. Lang, J. et al. Detecting and Quantifying Ataxia-Related Motor Impairments in Rodents Using Markerless Motion Tracking With Deep Neural Networks. in 2020 42nd Annual International Conference of the IEEE Engineering in Medicine Biology Society (EMBC) 3642–3648 (2020). doi:10.1109/EMBC44109.2020.9176701.

33. Schroer, R. J. et al. Natural History of Christianson Syndrome. Am. J. Med. Genet. A. 0, 2775–2783 (2010).

34. Koboldt, D. C. et al. VarScan 2: Somatic mutation and copy number alteration discovery in cancer by exome sequencing. Genome Res. 22, 568–576 (2012).

35. Gene Detail :: Allen Brain Atlas: Mouse Brain. https://mouse.brain-map.org/gene/show/88236.

36. Gilfillan, G. D. et al. SLC9A6 Mutations Cause X-Linked Mental Retardation, Microcephaly, Epilepsy, and Ataxia, a Phenotype Mimicking Angelman Syndrome. Am. J. Hum. Genet. 82, 1003–1010 (2008).

37. Ilie, A. et al. A Christianson syndrome-linked deletion mutation (Δ287ES288) in SLC9A6 disrupts recycling endosomal function and elicits neurodegeneration and cell death. Mol. Neurodegener. 11, (2016).

38. Ilie, A. et al. Assorted dysfunctions of endosomal alkali cation/proton exchanger SLC9A6 variants linked to Christianson syndrome. J. Biol. Chem. 295, 7075–7095 (2020).

39. Ohgaki, R. et al. The Na+/H+ Exchanger NHE6 in the Endosomal Recycling System Is Involved in the Development of Apical Bile Canalicular Surface Domains in HepG2 Cells. Mol. Biol. Cell 21, 1293–1304 (2010).

40. Nitta, K., Matsuzaki, Y., Konno, A. & Hirai, H. Minimal Purkinje Cell-Specific PCP2/L7 Promoter Virally Available for Rodents and Non-human Primates. Mol. Ther. Methods Clin. Dev. 6, 159–170 (2017).

41. Chan, K. Y. et al. Engineered AAVs for efficient noninvasive gene delivery to the central and peripheral nervous systems. Nat. Neurosci. 20, 1172–1179 (2017).

42. Hansen, S. T., Meera, P., Otis, T. S. & Pulst, S. M. Changes in Purkinje cell firing and gene expression precede behavioral pathology in a mouse model of SCA2. Hum. Mol. Genet. 22, 271–283 (2013).

43. Scoles, D. R. et al. Antisense oligonucleotide therapy for spinocerebellar ataxia type 2. Nature 544, 362– 366 (2017).

44. Pflieger, L. T. et al. Gene co-expression network analysis for identifying modules and functionally enriched pathways in SCA2. Hum. Mol. Genet. 26, 3069–3080 (2017).

45. Wu, Q.-W. & Kapfhammer, J. P. Modulation of Increased mGluR1 Signaling by RGS8 Protects Purkinje Cells From Dendritic Reduction and Could Be a Common Mechanism in Diverse Forms of Spinocerebellar Ataxia. Front. Cell Dev. Biol. 8, (2021).

46. Niewiadomska-Cimicka, A. et al. SCA7 Mouse Cerebellar Pathology Reveals Preferential Downregulation of Key Purkinje Cell-Identity Genes and Shared Disease Signature with SCA1 and SCA2. J. Neurosci. 41, 4910–4936 (2021).

47. Itoh, M., Odagiri, M., Abe, H. & Saitoh, O. RGS8 protein is distributed in dendrites and cell body of cerebellar Purkinje cell. Biochem. Biophys. Res. Commun. 287, 223–228 (2001).

48. Christianson, A. L. et al. X linked severe mental retardation, craniofacial dysmorphology, epilepsy, ophthalmoplegia, and cerebellar atrophy in a large South African kindred is localised to Xq24-q27. J. Med. Genet. 36, 759–766 (1999).

49. Brett, C. L., Wei, Y., Donowitz, M. & Rao, R. Human Na(+)/H(+) exchanger isoform 6 is found in recycling endosomes of cells, not in mitochondria. Am. J. Physiol. Cell Physiol. 282, C1031–1041 (2002).

50. Ouyang, Q. et al. Christianson syndrome protein NHE6 modulates TrkB endosomal signaling required for neuronal circuit development. Neuron 80, (2013).

51. Pescosolido, M. F., Ouyang, Q., Liu, J. S. & Morrow, E. M. Loss of Christianson Syndrome Na+/H+ Exchanger 6 (NHE6) Causes Abnormal Endosome Maturation and Trafficking Underlying Lysosome Dysfunction in Neurons. J. Neurosci. Off. J. Soc. Neurosci. 41, 9235–9256 (2021).

52. Zanni, G. et al. Mutation of plasma membrane Ca2+ ATPase isoform 3 in a family with X-linked congenital cerebellar ataxia impairs Ca2+ homeostasis. Proc. Natl. Acad. Sci. U. S. A. 109, 14514–14519 (2012).

53. OMIM Phenotypic Series - PS309510. https://omim.org/phenotypicSeries/PS309510?sort=geneSymbols.

54. Pescosolido, M. F. et al. Genetic and phenotypic diversity of NHE6 mutations in Christianson syndrome. Ann. Neurol. 76, 581–593 (2014).

55. Ouyang, Q. et al. Functional Assessment In Vivo of the Mouse Homolog of the Human Ala-9-Ser NHE6 Variant. eNeuro 6, (2019).

56. Gao, A. Y. L., Ilie, A., Chang, P. K. Y., Orlowski, J. & McKinney, R. A. A Christianson syndrome-linked deletion mutation (Δ287ES288) in SLC9A6 impairs hippocampal neuronal plasticity. Neurobiol. Dis. 130, 104490 (2019).

57. Prasad, H. & Rao, R. Amyloid clearance defect in ApoE4 astrocytes is reversed by epigenetic correction of endosomal pH. Proc. Natl. Acad. Sci. U. S. A. 115, E6640–E6649 (2018).

58. Kerner-Rossi, M., Gulinello, M., Walkley, S. & Dobrenis, K. Pathobiology of Christianson syndrome: Linking disrupted endosomal-lysosomal function with intellectual disability and sensory impairments. Neurobiol. Learn. Mem. 165, 106867 (2019).

59. Strømme, P. et al. X-linked Angelman-like syndrome caused by Slc9a6 knockout in mice exhibits evidence of endosomal–lysosomal dysfunction. Brain 134, 3369–3383 (2011).

60. Xu, M. et al. Mixed Neurodevelopmental and Neurodegenerative Pathology in Nhe6-Null Mouse Model of Christianson Syndrome. eNeuro 4, (2017).

61. Sikora, J., Leddy, J., Gulinello, M. & Walkley, S. U. X-linked Christianson syndrome: heterozygous female Slc9a6 knockout mice develop mosaic neuropathological changes and related behavioral abnormalities. Dis. Model. Mech. 9, 13–23 (2016).

62. Prasad, H. & Rao, R. The Na+/H+ Exchanger NHE6 Modulates Endosomal pH to Control Processing of Amyloid Precursor Protein in a Cell Culture Model of Alzheimer Disease. J. Biol. Chem. 290, 5311–5327 (2015).

63. Chakrabarty, P. et al. Capsid Serotype and Timing of Injection Determines AAV Transduction in the Neonatal Mice Brain. PLoS ONE 8, e67680 (2013).

64. Lukashchuk, V., Lewis, K. E., Coldicott, I., Grierson, A. J. & Azzouz, M. AAV9-mediated central nervous system–targeted gene delivery via cisterna magna route in mice. Mol. Ther. - Methods Clin. Dev. 3, (2016).

65. Ilie, A. et al. A potential gain-of-function variant of SLC9A6 leads to endosomal alkalinization and neuronal atrophy associated with Christianson Syndrome. Neurobiol. Dis. 121, 187–204 (2019).

66. Lin, Y.-T. et al. APOE4 Causes Widespread Molecular and Cellular Alterations Associated with Alzheimer’s Disease Phenotypes in Human iPSC-Derived Brain Cell Types. Neuron 98, 1141-1154.e7 (2018).

67. Corder, E. H. et al. Gene Dose of Apolipoprotein E Type 4 Allele and the Risk of Alzheimer’s Disease in Late Onset Families. Science 261, 921–923 (1993).

68. Strittmatter, W. J. et al. Apolipoprotein E: high-avidity binding to beta-amyloid and increased frequency of type 4 allele in late-onset familial Alzheimer disease. Proc. Natl. Acad. Sci. U. S. A. 90, 1977–1981 (1993).

69. Pohlkamp, T. et al. Endosomal Acidification by NHE6-depletion Corrects ApoE4-mediated Synaptic Impairments and Reduces Amyloid Plaque Load. 2021.03.22.436385 https://www.biorxiv.org/content/10.1101/2021.03.22.436385v1 (2021) doi:10.1101/2021.03.22.436385.

70. Shi, Y. et al. ApoE4 markedly exacerbates tau-mediated neurodegeneration in a mouse model of tauopathy. Nature 549, 523–527 (2017).

71. Garbern, J. Y. et al. A mutation affecting the sodium/proton exchanger, SLC9A6, causes mental retardation with tau deposition. Brain 133, 1391–1402 (2010).

72. Xu, M. et al. Mixed Neurodevelopmental and Neurodegenerative Pathology in Nhe6-Null Mouse Model of Christianson Syndrome. eNeuro 4, (2018).

73. Dong, J.-Y., Fan, P.-D. & Frizzell, R. A. Quantitative Analysis of the Packaging Capacity of Recombinant Adeno-Associated Virus. Hum. Gene Ther. 7, 2101–2112 (1996).

74. Love, M. I., Huber, W. & Anders, S. Moderated estimation of fold change and dispersion for RNA-seq data with DESeq2. Genome Biol. 15, 550 (2014).

75. Paul, S., Dansithong, W., Figueroa, K. P., Scoles, D. R. & Pulst, S. M. Staufen1 links RNA stress granules and autophagy in a model of neurodegeneration. Nat. Commun. 9, (2018).

76. Fowler, S. C. et al. A force-plate actometer for quantitating rodent behaviors: illustrative data on locomotion, rotation, spatial patterning, stereotypies, and tremor. J. Neurosci. Methods 107, 107–124 (2001).

